# Application of the *AMOCATI* R workflow to tumor transcriptomic data delineates the adverse effect of immune cell infiltration in immune-privileged organs

**DOI:** 10.1101/2024.06.18.596859

**Authors:** Paul Régnier, Nicolas Cagnard, Katrina Podsypanina, Guillaume Darrasse-Jèze

**Affiliations:** Immunology-Immunopathology-Immunotherapy (i3) Laboratory, INSERM UMR-S 959, Sorbonne Université, Paris, France; Biotherapy Unit (CIC-BTi), Inflammation-Immunopathology-Biotherapy Department (DHU i2B), Groupe Hospitalier Pitié-Salpêtrière, Assistance Publique-Hôpitaux de Paris (AP-HP), France; Structure Fédérative de Recherche (SFR) Necker, Université Paris Descartes, Paris France; Institut Necker Enfants Malades (INEM), INSERM U1151, CNRS UMR8253, Université Paris Cité, Paris, France; Université Paris Descartes, Sorbonne Paris Cité, Faculté de Médecine Paris Descartes, Paris, France

**Keywords:** Human cancers, Primary tumor biopsy, Transcriptome analysis, Immune cells infiltration, Immune-privileged organs, Overall survival, Algorithm, R language

## Abstract

Immune cells are present inside tumor tissue and can alter tumor growth. Expression profiles of human tumors hold transcripts from cancer cells and their microenvironment, including the infiltrating immune cells. Few standardized methods examine tumor immunobiology relying only on tumor transcriptome data. Using a new in-house developed R analysis workflow called *AMOCATI*, we classified 43 cancer types from 11,176 patients according to the degree of infiltration by 18 distinct immune cell subsets, measured by the abundance of their transcriptomic signature, and calculated its effect on the disease outcome. In about half of cancers affecting organs without immune privilege, immune cell infiltration has beneficial effects. In contrast, immune infiltration in cancers of immune-privileged organs (eye, testis and brain) confers poor prognosis. Moreover, transcriptional evidence of increased immune cell activity in immune-privileged cancer sites is associated with bad prognosis. Thus, our results suggest that the effect of immune infiltration may depend on the origin of the primary tumor.

**SIGNIFICANCE:** Our in-house developed computational R approach *AMOCATI* allows to easily download public transcriptomic and clinical data, classify and analyze them.

AMOCATI permitted us to define gene expression signatures associated with short- or long-term survival from 11,176 untreated patient unsorted biopsies in 43 types of cancer.

We present the level of infiltration of 18 types of immune cell subsets transcriptomic signatures and 50 immune-related pathways in all these cancers

Correlation between immune infiltration of the tumor and survival establishes a link between tumor tissue of origin and the overall effect of immune infiltration on survival.

Immune cell infiltration in tumors from ‘immune privileged organs’ correlate with shorter survival.

Contrary to what we observe in ‘hot’ tumors, biological pathways of immune response are associated with a short-term survival profile in these cancers.

## INTRODUCTION

The immune checkpoint blockade (ICB) therapies have revolutionized the treatment of cancer. It rests on the observation that anti-tumor immune cells may persist in the tumor at all stages of malignant disease but need to be properly disinhibited. The list of cancers that can be treated with immune checkpoint blockade (ICB) continues to grow. However, patients that can benefit from the ICB are often selected by trial and error. Predicting response is challenging because there are many populations of immune cells, and their behavior may be altered by the tumor microenvironment. Tumor-infiltrating immune cells have been extensively studied in mouse models(1), and several groups have linked immune infiltration to survival in individual human tumors types(2–4). To come up with prognostic biomarkers for selecting patients for ICB therapy, a number of reports used systems bioinformatics to look at the immune cell transcriptional signatures across multiple tumor types(5–11). The consensus is that, for a better prognostic power, the inflammatory biomarkers need to be considered in combination with information about clonal heterogeneity, somatic aberrations, gene copy number, presence of microRNAs, and activity of epigenetic processes in a given tumor. From a clinical standpoint, this amount of information is difficult to obtain before making a treatment decision. Thus, there is a need for a broad classification of tumor types into those that may or should not be treated with ICB.

Historically, immune infiltration has been ascribed widely opposing roles in the appearance and growth of cancer. Rudolph Virchow, the first to report that leukocytes frequently infiltrate tumors, thought that immune infiltrates facilitate cancer progression(12). Instead, Paul Erlich argued that the immune system protects the host from neoplastic diseases(13). Experimental evidence for anti-cancer immune surveillance, on one hand(14), and for pro-cancer inflammation(15) and immune tolerance(16–19), on the other, are mirrored by the complexity of the tumor-immune system interaction observed in the clinic. Tumor immune tolerance can be broken even at a late point of tumor development(20–24), and immune cell infiltration can be beneficial to patient survival(2, 25–28). But in some cases, immune infiltration is bad for patient survival(3, 29). In these instances, immune system may favor an immunoselection of more resilient tumor clones(1), or even favor tumor dissemination and metastasis(29). Efforts to classify tumors, patients and cancers according to their infiltrating immune cells abundance gave rise to the notion of “hot” and “cold” tumors, which are highly and poorly infiltrated by immune cells, respectively(30). However, multiple immune-related or tumor-related parameters are required at the moment of diagnosis to make therapeutic decisions for either a “hot” or a “cold” tumor(31).

For many tumor types, high immune cell density is a positive indicator of survival(32). In individual tumor types, the presence of a single immune subset can dominate the survival outcome. For instance, Zhang et al. showed that intra-tumoral CD3^+^ T cells were associated with increased survival in ovarian cancer(26). In breast cancer cohorts, Kohrt et al. showed that other immune subsets, such as CD4^+^ helper T lymphocytes, CD8^+^ cytotoxic T cells (CTLs) and CD1a^+^ DCs (DC2), are associated with increased survival(33). The NK cells infiltration in colorectal cancers (CRC) appears to have a positive role in disease control (directly, or by cooperation with DCs and CTLs), but its effect on survival is still being debated(34). Taking all of this into account, the relative proportions of two subsets of infiltrating immune cells can further refine survival analysis. Relative ratios of subsets critical for tumor growth, such as CTLs(35), DCs(36) or T_regs_(37), can affect prognosis differently, as shown for CD4^+^/CD8^+^ T cells ratio in colorectal cancer(38), or CD8^+^ T cells/T_regs_ ratio in ovarian cancer(39). However, for many tumor types, the relationship between immune infiltration and survival is not straightforward. Simultaneous evaluation of all immune system players in a tumor microenvironment may help retrieve prognostic information for cancer of any origin.

Analysis of transcriptomic and clinical data from public repositories, for example, TCGA, shows that gene expression can provide information about patient survival (40). The tumor transcriptomes also contain transcriptional signatures of various aspects of the immune response(41, 42). To retrieve these signatures, most published scripts, such as *CIBERSORT*(43), *EPIC* and others(44–46), rely on data deconvolution from bulk microarray or RNA-Seq source. One caveat of this approach is the requirement for an appropriate control to generate the deconvolution matrix. For maximal accuracy, controls are selected to match the tissue and the target cell population, to reproduce tissue-specific gene expression values. Such control data are rarely available when working on bulk transcriptomic datasets.

Here, we developed a novel computational method, called *AMOCATI* (*Algorithmic Meta-analysis Of Clinical And Transcriptomic Information*), which can be directly applied to the data from a prospective cohort, such as the TCGA one in this work. *AMOCATI* does not use deconvolution algorithms, but rather gene and pathway enrichments and correlations. As a standalone R package, *AMOCATI* workflow integrates functions to study the impact of individual genes or gene signatures on cancer patient survival. We used it to follow 18 immune cell types signatures and 50 immune-related pathways in 43 different cancer types in 11,176 patients from the Genomic Data Commons repository. By evaluating individual signatures and their combinations in the short- and long-term survivors classified with *AMOCATI*, we identified immune-system related transcriptional patterns that affect cancer outcomes.

## MATERIAL AND METHODS

### Generation of immune cell population-specific gene signatures

We generated immune cell subset-specific gene signatures using publicly available GEO transcriptomic datasets (**Table S1, *GEO_Datasets_Signatures* sheet**) and cross-validated them. After downloading raw signatures specific for cell subsets, we manually refined them using successive heatmaps and Venn diagrams in order to remove as many of low-expressed and poorly specific genes as possible. We defined 18 distinct gene signatures, each one as specific as possible for its related immune cell subset. To illustrate their specificity, we created four Venn diagrams compiling several signatures together, as displayed in the **Figure S2D**. In the top-left diagram, we compared signatures from seven myeloid cell subsets (pDCs, cDC1, cDC2, cMo and ncMo) and in the bottom-left we included signatures from five lymphoid cell subsets (T_regs_, T_effs_, B, NK and CTL). The naïve and activated/memory subsets of T_regs_ and T_effs_ are also shown in the top-right and bottom-right diagrams.

### *All-in-one* data file generation

This *AMOCATI*-specific file format was created to compile all transcriptomic and clinical data about one cancer dataset. Each row contains information about one patient. The first column *CaseUUID* is a unique patient identifier. The second column *vitalStatus* contains the current vital status of a patient (*Alive* or *Dead*). The third column *survivedDays* contains the number of days survived from tumor diagnosis to the last patient clinical follow-up. Finally, all subsequent columns contain information on absolute expression of about 60,000 gene transcripts as determined by RNA-Seq (one per column) with the name of each column uniquely identifying each gene (using HGNC identifiers). In the present study, the end of the survival analysis was fixed to five years (1,825 days), meaning that 1) every patient alive after this cut-off has his survival days fixed at 1,825 and 2) every patient who died after this cut-off is considered alive for the study. The 1,825 days cut-off is useful to the further *metaResults* generation algorithm: this prevents inconsistencies in calculations because of the very-long surviving patients in one or more groups.

### *MetaResults* algorithm details

This two-step method measures impact of each individual gene on patient survival within a given cancer cohort. As the survival time after diagnosis is one of the few clinical parameters included in both TCGA, CGCI and the TARGET datasets, we focused on this value to create the survival curves. Other clinical parameters, such as tumor grade, age, or gender, are indicated only for a fraction of samples, thus limiting the overall group sizes.

In the first step, patients whom *survivalDays* field is empty or equal to zero are removed from the analysis. Then, the algorithm performs a bootstrap operation on a random sample of 50% of the remaining cohort. Within an iteration *i*, the algorithm computes several statistics: µ_gi_ which represents the mean of gene g expression and SD_gi_ which represents the standard deviation of gene *g* expression. It also creates survival curves for patients with low-, intermediate- and high-expression using fixed 15% as low- and high-cut-off). From these, the algorithm computes the log-rank test statistic and p-value for “intermediate”/“low” (respectively S-IL_gi_ and p-IL_gi_) and “intermediate”/“high” (respectively S-IH_gi_ and p-IH_gi_) group comparisons. Each of the previously mentioned values is stored into a text file to be used in the next part of the algorithm. The same instruction is independently iterated 96 times.

In the second step, the algorithm integrates results obtained from the 96 iterations for each gene to estimate the robustness and reliability of the obtained metrics. For both “intermediate”/“low” and “intermediate”/“high” groups comparisons, it calculates the mean and SD of the log-rank test statistics (µ[S-IL_g_], SD[S-IL_g_], µ[S-IH_g_] and SD[S-IH_g_]), and the mean of the log-rank test p-values (respectively µ[p-IL_g_] and µ[p-IH_g_]). These values are used to compute the SNR[IL_g_], defined as µ[S-IL_g_] divided by SD[S-IL_g_] for “intermediate”/“low” comparison and the SNR[IH_g_], defined as µ[S-IH_g_] divided by SD[S-IH_g_] for “intermediate”/“high” comparison. The two SNR values are then weighted by the difference between 1 and the respective mean of log-rank tests p-values (µ[p-IL_g_] or µ[p-IH_g_]) and finally added together to form the final Gene Score GS_g_. The means of every y g-related µ_gi_ and SD_gi_ are computed (respectively µ[µ_g_] and µ[SD_g_]) and used to calculate the global gene g SNR (SNR[G_g_]) defined as µ[µ_g_] divided by µ[SD_g_]. In parallel, the full patients survival curves are constructed using the same fixed 15% threshold for low- and high-expressing gene g patients, allowing to classify the overall gene effect on survival into one of the four predefined settings: *Correlation* (the more the gene is expressed, the better is the survival), *Inverse Correlation* (the more the gene is expressed, the worse is the survival), *Paradox* (low and high expression of the gene leads to better survival compared to intermediate expression) and *Inverse Paradox* (low and high expression of the gene leads to worse survival compared to intermediate expression). The resulting setting is called S_g_. Finally, GS_g_, SNR[G_g_] and S_g_ values are exported for each gene and constitute the main output values of the *metaResults* file, complete with the gene name and the cut-off percentile (15% here). A full detailed graphical version of the algorithm is presented as UML diagrams in **Figure S1B** and **Figure S1C**.

The GS_g_ reflects the strength and the robustness for the gene g to separate low-, intermediate- and high-expressing patients respective survival curves. The higher the score, the stronger is the effect on survival, and the less important is the variation upon random iterations. The SNR[G_g_] reflects the global gene g variation of expression between the iterations. The higher this metric, the more expressed the gene and the less is the variation between patients. The setting S_g_ represents the pre- defined category in which the gene g was classified according to its overall effect on low-, intermediate- and high-expressing patients survival curves. Of note, threshold values for GS_g_ used to isolate genes that will compose the final classification signatures typically start from two but can go higher if needed. On the contrary, SNR[G_g_] threshold was set to two for all the analyzed cancers.

### *Clinical Score* algorithm details

This algorithm is used to compute a *Clinical Score* (CS) for each given patient and gene signature. This CS represents the propensity of a signature to be expressed in a manner that is favorable to a patient survival: the higher, the better for the survival, and *vice versa*. To explain the algorithm, let’s take a hypothetical gene signature composed of 2 genes, A and B. In the *metaResults*, gene A was classified as *Correlation* and B as *Inverse Correlation*. To calculate the CS, the algorithm takes the first gene of the signature, A, and looks in the associated *metaResults* table the setting for this gene, here *Correlation*. Using the fixed cut-off percentile (set here to 15%), the algorithm computes the associated gene A “intermediate”/“low” and “intermediate”/“high” expression cut-offs in order to separate the patients into these three groups, as previously mentioned. Then, for every patient p, the algorithm looks at the expression of the gene A (E_pA_) and decides to increment or decrement the score by E_pA_ given the verity table presented in the **Figure S2**. In our example, if a patient is expressing the gene A as “high”, this is favorable to his survival, so the CS is incremented by E_pA_. If another patient expresses the gene A as “low”, this is not favorable to his survival, so the CS is decremented by E_pA_, which actually represents the subscore for the current gene A. And so on for every patient. Then the sequence is repeated for gene B. Because its setting is *Inverse Correlation*, if a patient is expressing the gene B as “high”, this is bad for his survival, so the CS is decremented by E_pA_. If another patient expresses the gene B as “low”, this is now favorable to his survival, so the CS is incremented by E_pA_. Then, the gene subscores for each patient are summed up. Finally, the algorithm divides each patient’s total CS by the number of genes composing the signature, making the final CS an average of the individual gene subscores for each patient.

### Custom gene sets enrichment analysis

We decided to focus here on several publicly available gene set databases: Gene Ontology, KEGG, PANTHER, Reactome and WikiPathways which are constantly kept up-to-date and contain a plethora of information about pathways and biological functions. To be sure to get the most recent gene sets from these databases, we wrote R functions that directly contact the dedicated servers, allowing us to construct our gene sets database that we can fully update at any time on our own. The whole gene sets and their gene contents are then stored in an offline text file. The final gene sets file we generated and used throughout the manuscript was composed of 5,077 gene sets, as recent as March 2021. This way, we were able to avoid using other developed tools (like *GSEA* or other web-based tools) that mainly use outdated versions of the cited databases. The pathways enrichment in patients was assessed using the *GSVA* R package, following the authors instructions. Next, statistical significance between our groups of interest (STS and LTS here) was assessed using standard *limma* R package, following authors instructions.

### Generation of network graphs and meta-clustering of pathways

Among the enriched pathways, we only kept the ones that were significantly enriched (adjusted p-value < 0.05) and for which the enrichment statistic absolute value was above a defined threshold (usually 0.05 or 0.1). Next, we computed the similarity matrix, which contains pairwise similarity metrics for every retained pathway, defined here as the percentage of common genes between two gene sets and used this matrix within the *igraph* R package to create the final network graphs. On a network, each node represents a given gene set, which can be connected if the similarity value between two of them is greater than a defined similarity threshold (typically around 0.2). Next, we removed the nodes which were not connected to any other one as well as clusters containing too few pathways (usually between 3 and 5). Finally, we determined clusters of pathways around highly connected and dense node clouds and extracted their contents in gene sets, using dedicated functions in the *igraph* R package. We then used the information to manually annotate every cluster for each selected cancer, leading to the graphs presented in the **Figure 2**.

**Figure 1.**
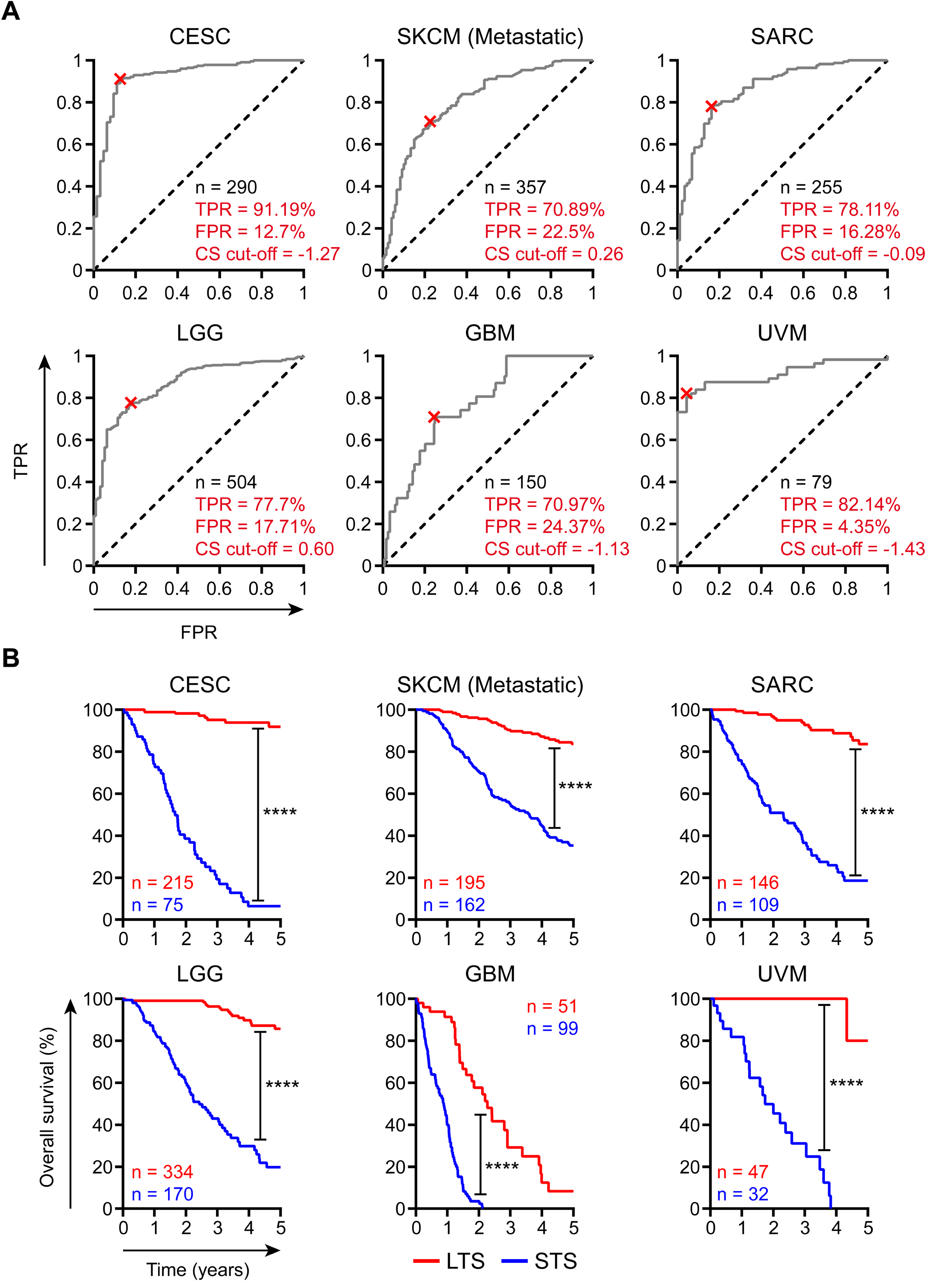
*Clinical Score* (CS) computation algorithm determines transcriptomic gene signatures discriminating long-term and short-term survivors in six cancers. (A) ROC curves showing true and false positive rates (TPR and FPR) together with optimal CS cut-off obtained by the determination of Youden Index point coordinates in six selected cancer datasets (SKCM (Metastatic), CESC, SARC, GBM, LGG, UVM). CS values for each patient were obtained using the CS algorithm presented in **Figure S1D**. **(B)** Survival curves showing the STS and LTS separation in six cancer datasets obtained using the previously determined optimal CS cut-off.

**Figure 2.**
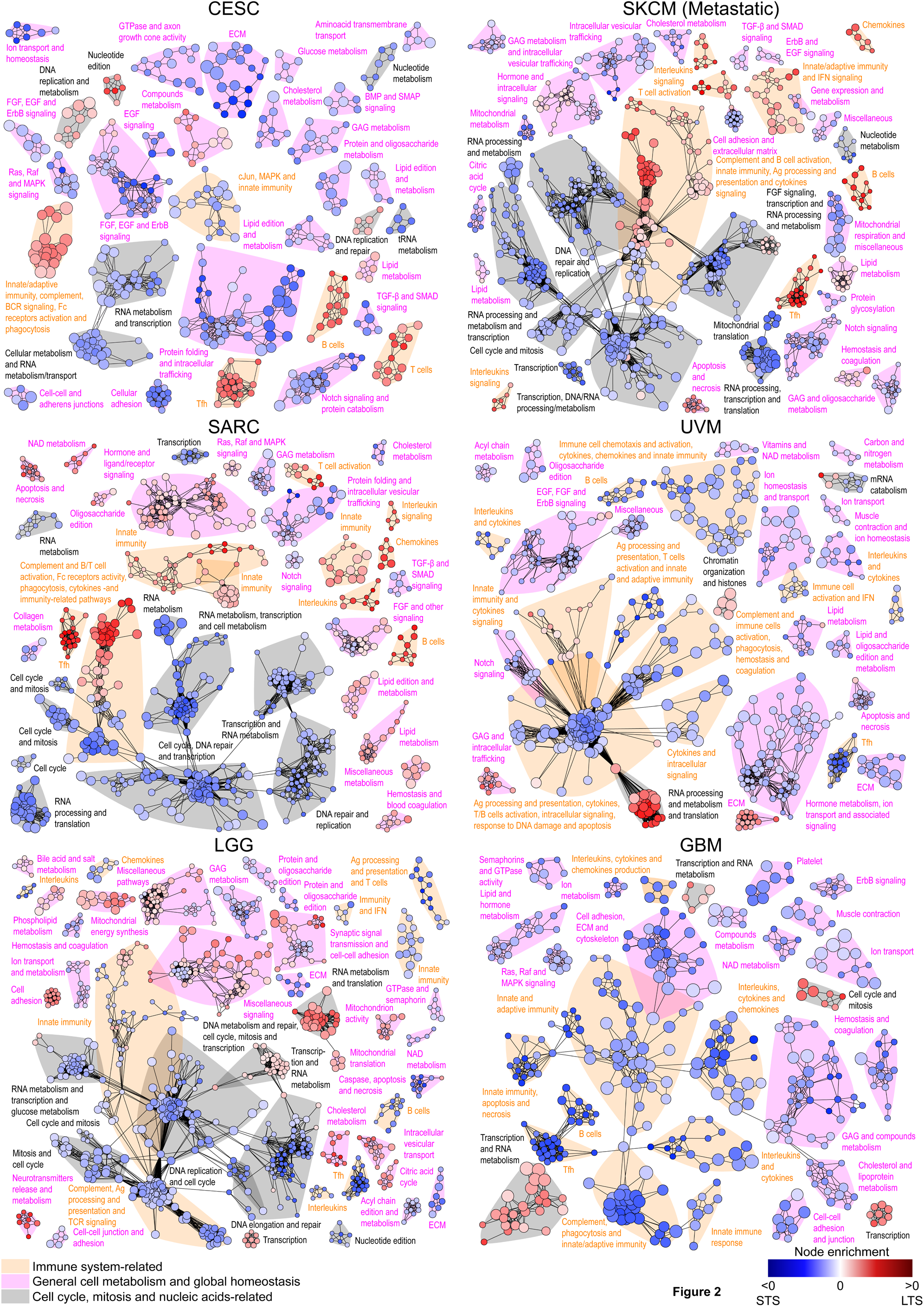
Enrichment of immunological functions among biological pathways distinguishing LTS and STS groups in six selected cancers. Network representation of pathway enrichment analysis from the list of pathways differentially expressed between LTS and STS for each depicted cancer. Two thresholds were used to construct each graph: the p-value threshold to select the significant individual nodes (pathways), and the similarity (percentage of identical genes between two pathways) threshold to connect the remaining nodes together. A color scale is applied to each node, using for each pathway its computed enrichment as metric. Negative values (more enriched in STS) increasingly go from blue to white, whereas positive values (more enriched in LTS) increasingly go from white to red (bottom right legend). A second discrete color scheme is used, allowing to categorize each cluster found during graph construction. Immune system-related pathways are highlighted in orange, general cell metabolism and global homeostasis are in purple and cell cycle, mitosis and nucleic-acids-related are in grey (bottom left legend).

### Receiver Operating Characteristic analysis and Kaplan-Meyer survival curves

We used *ROCit* R package to construct ROC curves based on the CS values computed for the classification signature in each depicted cancer. Then, the Youden Index point coordinates were computed and used to determine the optimal CS cut-off value to use to best distinguish patients according to their actual survival state (*Dead* or *Alive*). This led to the generation of two distinct groups of patients: the ones who have a low CS value (below the threshold) and subsequently labelled as Short-Term Survivors (STS), and the ones who have a high CS value (above the threshold) and subsequently labelled as Long-Term Survivors (LTS). Finally, we constructed the Kaplan-Meyer survival curves using this separation between STS and LTS groups with the *survival* R package.

### Statistical analysis

Differences between survival curves were assessed using log-rank tests. Differential analyses of pathway enrichments were assessed using moderated t-tests included in the *limma* R package. Correlation analyses were performed using Pearson correlation coefficient with associated default inference test within R base functions. P-values < 0.05 were deemed significant.

### Software availability

*AMOCATI* source code is available as a standalone R package which is fully accessible on the following GitHub repository: https://github.com/PaulRegnier/AMOCATI. Further information about installation is provided on the same GitHub link. Additionally, a tutorial showing an example analysis made with *AMOCATI* is available on the following URL: https://paul-regnier.fr/tutoriel-amocati/.

## RESULTS

### Development of an algorithmic tool for meta-analysis of clinical and transcriptomic data

Our first objective was to design a workflow for easy analysis of TCGA, TARGET and CGCI cancer projects coming from GDC repository. **Table S1, *Cancer Datasets* sheet** summarizes the datasets used in this work and the associated version number of the data. As shown in **Figure S1A**, the first function of *AMOCATI* is to download all RNA-Seq and Clinical data for a given cancer, normalize and pool them in a single data text file (called *all-in-one* file, see the **Material and Methods** section).

Another feature of *AMOCATI* is to measure the impact of each gene on patient survival. This is made in two separate steps, which are fully detailed in the **Material and Methods** section. At the end of this process, these algorithms create an output table called *metaResults* (MR), containing, for each gene, its estimated impact on patient survival in a given cancer, as well as metrics estimating strength and likeliness/robustness of the impact. First, for each gene independently, *AMOCATI* applies a bootstrapping method. This involves randomly sampling half of the cohort for a given number of independent iterations (96 here) and using it to 1) construct the associated survival curves for patients with low, intermediate and high expression for the gene and 2) compute several metrics for further analysis (**Figure S1B**). In the present work, we used 15% as a cut-off for low- and high-expressing groups, thus assigning 70% of patients in the intermediate-expressing group.

The second step integrates the results obtained in the 96 independent iterations for each gene (**Figure S1C**). The main aim is to compute a Gene Score (GS), which estimates the impact of a gene expression on patient survival and its performance upon iterations (**Figure S1C**, red and blue paths). A high GS_g_ means that a transcript has a strong and specific impact (positive or negative) on survival.

Along with GS_g_, the second step returns two other metrics for each gene. The first one is a global signal-to-noise ratio, SNR[G_g_]. It estimates the overall expression strength relative to background noise independently of the different expression groups (“low”, “intermediate” or “high”) or the survival time information (**Figure S1C**, green path). This metric ultimately allows to eliminate genes which are poorly expressed and/or show hyper variation upon bootstrapping iterations. We used a threshold of 2 for the SNR[G_g_]. The second metric, called the setting S_g_, is the overall gene effect on survival for every patient from the cohort, which is independent of the bootstrapping iterative method (**Figure S1C**, purple path). This metric is assigned one of the following pre-defined settings: *Correlation*, *Inverse Correlation*, *Paradox* or *Inverse Paradox*.

To summarize, the output MR table contains the following information for each gene: its name, its cut-off value c (here 15% for every gene and dataset), its SNR[G_g_], its GS_g_ and its setting S_g_. Full details are available within the **Material and Methods** section.

### AMOCATI can be used to generate a classification signature that efficiently separates long- and short-term survivors

To retrieve genes for a classification signature, we plot for each gene g its SNR[G_g_] versus its GS_g_ in a single dot-plot. As an example, **Figure S2A** shows the thresholds used to construct a classification signature within the BRCA dataset. The 99 genes contained in the upper-right quadrant all have GS_g_ and SNR[G_g_] values above 2.25 and 2 respectively, and thus are the most likely to robustly separate the patients into long-term survivors (LTS) and short-term survivors (STS). Thus, *AMOCATI* returns 99 genes in the BRCA classification signature (**Figure S2A**).

*AMOCATI* also works with any gene signature and computes two scores for each patient and each gene signature. The *Quantitative Score* (QS) is an estimator of the gene signature abundance and is defined as the arithmetic mean of all the gene expressions contained in the signature. The *Clinical Score* (CS) (**Figure S1D** and **Material and Methods** section) is used to estimate whether a gene (CS_g_) or a gene signature (CS_gs_) is associated with survival or death in a patient cohort. The QS and CS may be compared only between patients or cohorts, and not between signatures.

Then, the CS_g_ is computed for every gene of the signature in each patient, and the mean of the CSs for the signature constitutes the CS_gs_, as described in **Material and Methods** section. To compute the CS, the algorithm separates patients in two groups according to the CS_gs_ values: STS who have the lowest CS_gs_ values and LTS who have the highest CS_gs_ values. We applied this method on patients of six selected cancers (CESC, SKCM (Metastatic), SARC, LGG, GBM and UVM). First, we computed their CS_gs_ for the associated classification signature (summarized for these cancers in the **Table S1, *Classification Signatures* sheet**). We then used Receiver Operating Characteristic (ROC) curves together with Youden Index point determination to select an optimal cut-off for the CS value (**Figure 1A**). This allowed us to allocate patients between STS (CS below the cut-off) and LTS (CS above the cut-off) groups according to each patient CS_gs_ value, which finally permitted the construction of their respective survival curves (**Figure 1B**). We observed a very strong and significant separation of LTS and STS in the selected cancers. In all cases, actual True Positive Rates (TPR) were > 70% and False Positive Rates (FPR) were < 25%. Similar or even better results were also found in all other cancers (data not shown). Thus, *AMOCATI* method can identify exceptional cases in a cohort of cancer patients using only transcriptional data and time of survival.

To evaluate the relative performance of our method, we calculated CS_gs_ for gene signatures that were previously reported to have an impact on breast cancer survival(47) as well as 10 random gene signatures. We calculated CS_gs_ associated with each of these signatures and then applied *AMOCATI* to separate LTS and STS. We then extracted the χ^2^ values associated with the log-rank test done on survival curves separating LTS and STS. The **Figure S2B** shows that in BRCA cancer dataset, the *AMOCATI*-generated signature is the best one to detect a difference between LTS and STS, superior to other breast cancer related (grey) and unrelated/random (blue) signatures. In addition, the absolute difference of Restricted Mean Survival Time (RMST) between STS and LTS survival curves was the highest when *AMOCATI*-generated classification signature was used. It is important to note that *AMOCATI*-generated classification signatures were generated using GDC Portal cancer database, which is not the case for the gene signatures collected from the literature.

To determine if *AMOCATI* method allows to establish classification signatures that may predict survival on an unknown dataset, we used the BRCA dataset and split it into two subdatasets A and B of equivalent size using four in-a-row random shuffling. For each subdataset, we generated the MR_A_ and MR_B_ tables and defined two classification signatures (Sign_A_ and Sign_B_). Then, we applied Sign_A_ and Sign_B_ on each subdataset (A and B), using its corresponding MR (MR_A_ or MR_B_, respectively) file to apply the signatures. The survival curves separating LTS and STS using matching and non-matching signatures show that efficient separation of LTS and STS can be only achieved when applying Sign_A_ or Sign_B_ on the matching subdataset (**Figure S2C**). This result implies that *AMOCATI* effectively identifies exceptional cases with respect to survival in a given patient cohort, which makes it an efficient descriptive tool, but also underlines that *AMOCATI* cannot be used as a prognostic nor a predictive tool.

In summary, the *AMOCATI* algorithms use only transcriptional data and length of survival to turn out simple indexes that estimate 1) the qualitative and quantitative impact of any gene on survival, 2) the relative abundance of any gene signature in a given dataset and 3) the relative impact of any gene signature on patient survival. Most importantly, we show that our classification signatures allow to define exceptionally good and exceptionally poor survivors (LTS and STS, respectively) in any given group of patients.

### Pathway enrichment analysis reveals opposite effects of immune system infiltration on patient survival depending on the cancer type

We reasoned that the transcriptional profiles of the STS and LTS groups classified by *AMOCATI* may contain information about the biological underpinnings of their different fates. To test this, we performed pathway enrichment analysis using gene sets directly recovered from Gene Ontology, KEGG, Reactome, PANTHER and WikiPathways databases and created network graphs involving the most differentially regulated pathways between LTS and STS in the same six cancers. We found that immune system-related pathways and clusters (in orange) were enriched in each of these cancers, but in different outcome groups (**Figure 2**).

For CESC and SKCM (Metastatic) (top graphs), as well as SARC (middle-left graph) cancers, the immune-related pathways were associated with survival, as they were enriched in LTS. We found common clusters between these three cancers, such as innate immunity, complement activation, immune system-related signaling pathways and adaptive immune response via T/B cells. Those results suggest that in CESC, SKCM (Metastatic) and SARC cancers, immune cell infiltration and canonical immune system processes are associated with good prognosis.

In UVM (middle-right graph), LGG and GBM (bottom graphs), immune-related pathways were enriched in STS, including clusters for innate and adaptive immunity, complement activation, cytokine/chemokine secretion/production and immune cells migration/chemotaxis. These results show that the immune cells infiltration and general immune system processes are associated with poor survival in LGG, GBM and UVM patients. Of note, the immune-related pathways and clusters are broadly shared between all the six cancers, indicating that the same immune function may have a good or a bad impact on survival depending on the origin of a tumor.

We conclude that the unsupervised analysis of gene set networks in the STS and LTS groups classified by *AMOCATI*-generated gene signatures shows evidence of immune system involvement in the control of cancer outcomes. However, the effect on survival can be positive or negative, dependent on the cancer type and location.

### In tumors from immune-privileged sites, transcriptional evidence of infiltration with individual immune cell subsets is associated with negative effect on survival

We sought to investigate whether the transcriptional evidence of infiltration with a certain immune cell subset was related to the overall effect on survival. We selected 18 immune cell subsets and computed their individual gene signatures based on the previously published works (**Table S1, blue-, green-, red- and orange-colored sheets** therein). The resulting subset-specific signatures have very few overlaps between them (**Figure S2D**). We used *AMOCATI* to compute the QS for each subset and each patient and plotted the mean QS values for each subset in a row-normalized heatmap to compare different tumor types (**Figure 3A**). We saw that cancers display a very broad range of QS: some cancers are infiltrated with low amounts of immune cells (including UVM and SARC), whereas others are highly infiltrated with immune cells (like KIRC, STAD, HTMCP-CC, ESCA, BRCA, LUAD, and LUSC). However, these groups did not correspond to those determined by the unsupervised pathway enrichment analysis. Indeed, many cancer types (including GBM, LGG, CESC and SKCM(Metastatic)) display high abundance for some cell subsets and low abundance for others, suggesting that outcomes may depend on infiltration by a specific subset only.

**Figure 3.**
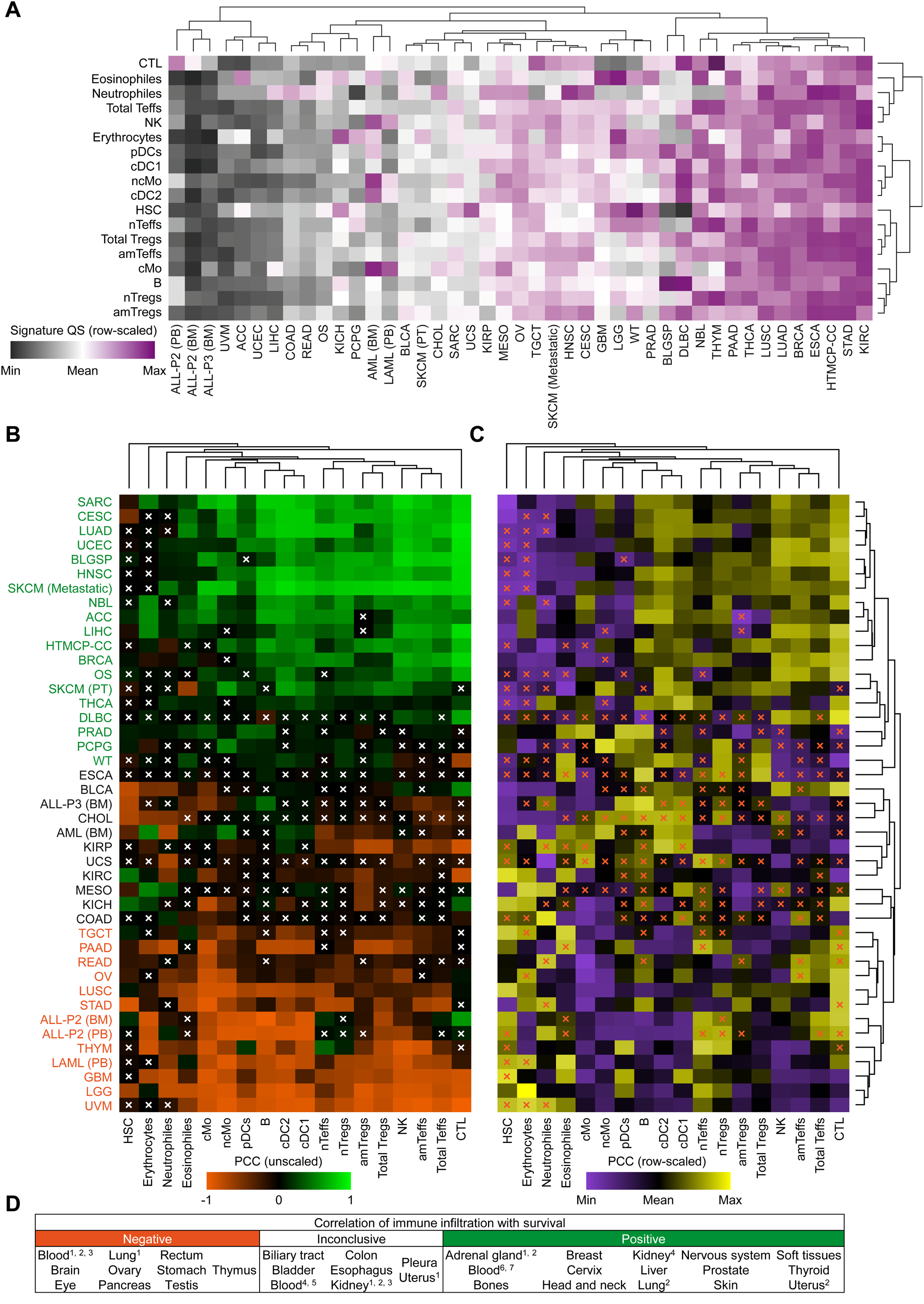
Quantification and correlation with overall patient survival of 18 cell subsets transcriptomic signatures in 43 cancers reveals negative impact of immune cell infiltration in tumors from immune-privileged sites. (A) Row-scaled heatmap showing QS computed for each cell signature in each cancer. Unsupervised clustering analysis separates low-infiltrated (grey) from high-infiltrated (pink) tumors. **(B)** Heatmap showing the Pearson’s coefficient of correlation (PCC) between QS and CS reflecting positive or negative correlation to survival of each cell signature in each cancer. An orange-black-green global color scale is used going from -1 to +1, according to each PCC value. Dendrogram clustering were then computed on both rows and columns of the heatmap. White crosses mark non-significant PCC values (p-value > 0.05). **(C)** Heatmap showing the same data as in **(B)** but using a row-normalized color scale, going from the minimum of each row (purple) to the maximum of each row (yellow) with a black middle color, in order to better dissect every cell population impact on survival in a given cancer. Dendrogram clustering were then computed on both rows and columns of the heatmap, giving the same clustering as in **(B)**. Orange crosses mark non-significant PCC values (p-value > 0.05). **(D)** Table summarizing the cancers classification according to the overall impact of their respective immune cells infiltration.

To test whether some of the cell subsets were individually associated with survival, we first computed the CS of every cell population signature in each patient and cancer. Then, we computed the Pearson’s coefficient of correlation (PCC) between QS (abundance) of each cell population signature and the CS (outcome) in the entire cohort and plotted the PCCs for each cancer on a single heatmap (**Figure 3B**). QS/CS correlations identified cancers in which immune infiltration is broadly associated with survival (green), and those in which any immune infiltration shortened survival (orange). Importantly, almost every immune cell subset signature in GBM, LGG, TGCT and UVM types of cancer shows an adverse effect on patient survival. This indicates that despite the variety in the abundance of individual subsets, any immune infiltration is harmful for patients with cancers originating from immune-privileged sites.

To better delineate which cell populations dominate survival outcomes in a single cancer, we changed the color scaling method used in the heatmap. The **Figure 3C** presents the same results as in **Figure 3B**, except that the colors are now row-normalized. Rows cannot be directly compared together anymore, but it allows to visualize cell populations that are deleterious (purple) or beneficial (yellow) for survival in a given tumor type. For instance, in cancers in which immune infiltration is beneficial to a patient’s outcome (green), the presence of cDC1 and cDC2 (but not pDC) populations is associated with high CS, as well as that of NKs and amT_effs_, while amT_regs_ (*am* standing for activated/memory) are only weakly associated with survival. It suggests than in these cancers, immune effector cells (represented by amT_effs_, NK cells and CTL) and cDCs are at least partially responsible for better disease outcomes. In contrast, in the majority of cancers in which immune infiltration was associated with poor survival (orange color in **Figure 3B**), and notably in tumors of the immune-privileged sites of the eye, brain and testis (UVM, LGG, GBM and TGCT), we observed that even the known anti-cancer players (NK cells, cMo, ncMo, cDC1, cDC2 and B cells) are not associated with high CS. Intriguingly, this comparison shows that the presence of naïve T cells (nT_regs_ and nT_effs_) is a relatively positive sign for these patients, despite the fact that they show negative correlation with survival in cross-cancer comparisons.

Taken together, the outcomes of tumor immune infiltration (**Figure 3B** and **Figure 3C**) and the overall quantitative levels of infiltration (**Figure 3A**) show that the extent of immune infiltration does not always correlate with survival: some cancers are highly infiltrated and this infiltration is associated with good prognosis (like BRCA or LUAD); other cancers are poorly infiltrated, but the lack of infiltration is associated with good prognosis (like UVM or READ); third kind is highly infiltrated, but the infiltration confers bad prognosis (like LUSC or STAD); and the last kind is poorly infiltrated, but should better be (like ACC or UCEC).

We conclude that tumor transcriptomic data may be used to estimate and compare the extent of immune infiltration in different cancers and its impact on survival (summarized in **Figure 3D**). The effect of the immune infiltration is specific to each cancer type but was reproducibly detrimental in cancers of immune-privileged organs (UVM, LGG, GBM, OVM, TCGT).

### Transcriptional evidence of increased immune cell activity in immune-privileged cancer sites is associated with bad prognosis

To validate the results of the pathway enrichment analysis, we studied the correlation between the abundance (QS) of the 18 cell subset signatures as previously defined, and the QS of immune-related biological pathways that are differentially regulated between STS and LTS. We next isolated 50 representative pathways that were commonly and significantly (adjusted p-value < 0.05) differentially regulated between STS and LTS in the six previously selected cancers. For each cancer, we computed the PCCs between the QS of each of the 18 subset signatures and the QS of each of the 50 immune-related pathways, in STS and in LTS. The resulting correlations were split in two different heatmaps, one for STS (left) and the other for LTS (right). (**Figure 4A** for SKCM (Metastatic), **Figure 4B** for UVM, **Figure S3A** for SARC, **Figure S3B** for LGG, **Figure S3C** for CESC and **Figure S3D** for GBM). In STS group of SKCM (Metastatic) (**Figure 4A, left**) the correlations across cell types and pathways rarely reach 0.9. In contrast, in the LTS these correlations tend to go closer to 1 (**Figure 4A, right**). The same relationship is observed in SARC (**Figure S3A**) and CESC (**Figure S3C**). A strong correlation between the transcriptional evidence of the immune-related pathways and the transcriptional evidence of infiltration by different immune cell populations in LTS shows that in cancers with beneficial effects of immune infiltration, a high immune cell activity is required for a positive effect on survival.

**Figure 4.**
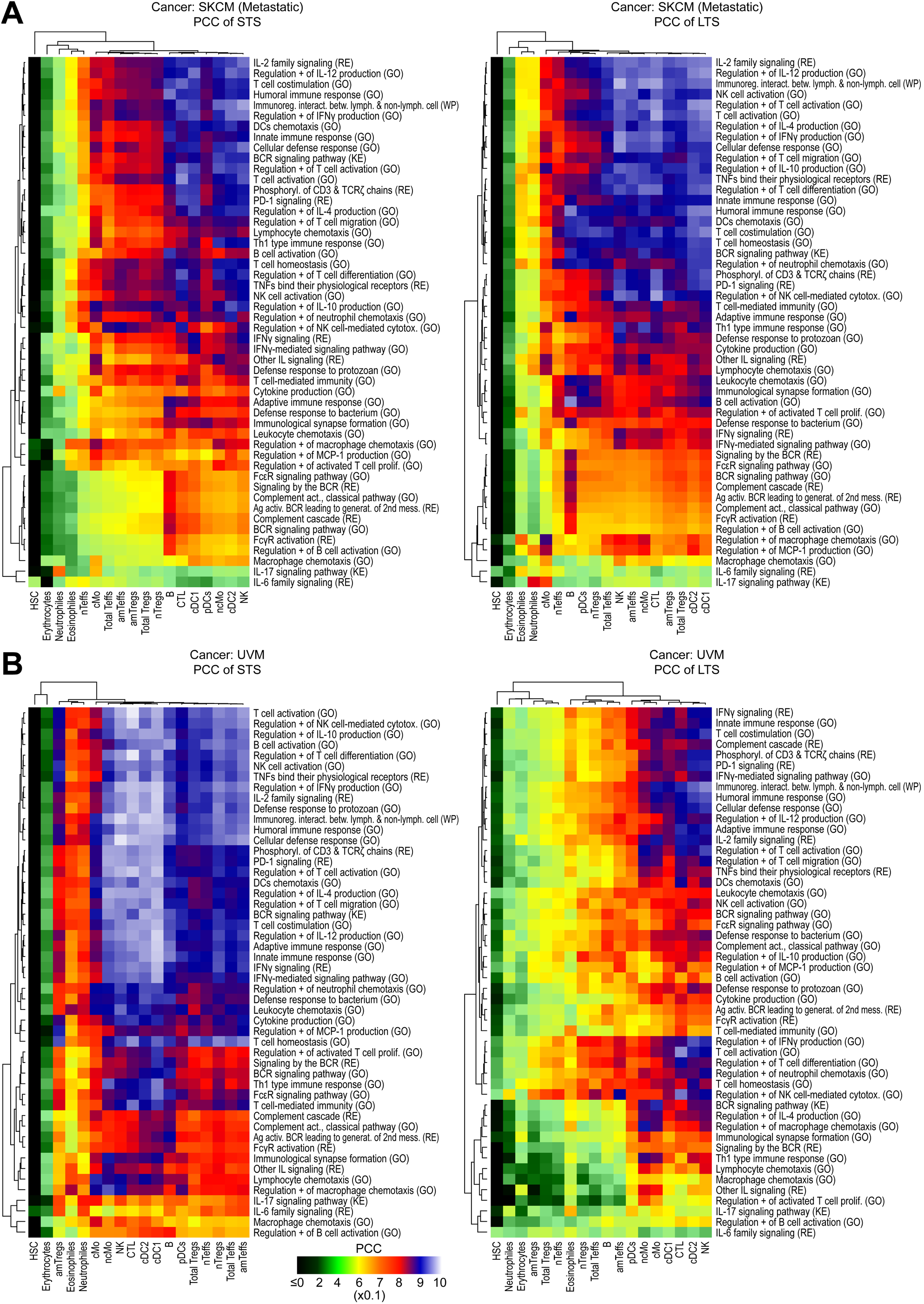
Correlations between functional pathways and cell signatures QS are altered in UVM as compared to SKCM (Metastatic), evoking an immunopathological reprogramming of infiltrating cells in immune-privileged cancer sites. Heatmap representations of the PCCs computed between cell populations gene signatures QS and selected immune system-related biological pathways QS in the SKCM (Metastatic) cancer **(A)** and the UVM cancer **(B)**. The 50 pathways that are represented are commonly and significantly dysregulated between LTS and STS of six selected cancers (SKCM (Metastatic), CESC, SARC, UVM, LGG and GBM) and are restricted to the immune system. In each case, STS and LTS patients are plotted separately in different heatmaps (left for STS and right for LTS). Dendrogram clustering were realized for both rows and columns. Black to yellow colors represent PCCs between 0 and 0.6, yellow to orange between 0.6 and 0.7, orange to red between 0.7 and 0.8, red to blue between 0.8 and 0.9, and blue to white between 0.9 and 1.

An opposite relationship is observed when we look at UVM cancer (**Figure 4B**), a cancer of an immune-privileged site: high correlations between immune pathways and populations are present in STS (**Figure 4B, left**), whereas they almost totally disappear in LTS (**Figure 4B, right**), for instance with the *T cell co-stimulation*, *Regulation + of T cell activation* and *T cell activation* pathways. This phenomenon is also observed in other immune-privileged cancers such as LGG (**Figure S3B**) and GBM (**Figure S3D**). Thus, an increased immune cell activity accompanies immune infiltration both in tumors with beneficial and those with deleterious effects of immune infiltration.

Importantly, the evidence of co-regulation between subset and pathway transcripts held for almost every individual pathway/subset pair, regardless of the tumor type. To look at individual pairs, we subtracted the PCCs obtained for LTS and STS for each immune pathway and cell population, then plotted the subsequent results in a single heatmap for each cancer. For SKCM (Metastatic) we observed that the majority of differences of correlations are greater than zero (red), thus, more strongly correlated in LTS than in STS (**Figure 5A, left**). Similar relationship is observed in SARC (**Figure S3E**) and CESC (**Figure S3F**). On the other hand, in UVM almost all differences of correlations are less than zero (blue), thus, more strongly correlated in STS than in LTS (**Figure 5A, right**). We observed the same pattern in other immune-privileged cancers such as LGG (**Figure S3G**) and GBM (**Figure S3H**). Thus, the positive or negative effect of tumor-immune system interaction on survival is conserved between individual pathway/subset pairs.

**Figure 5.**
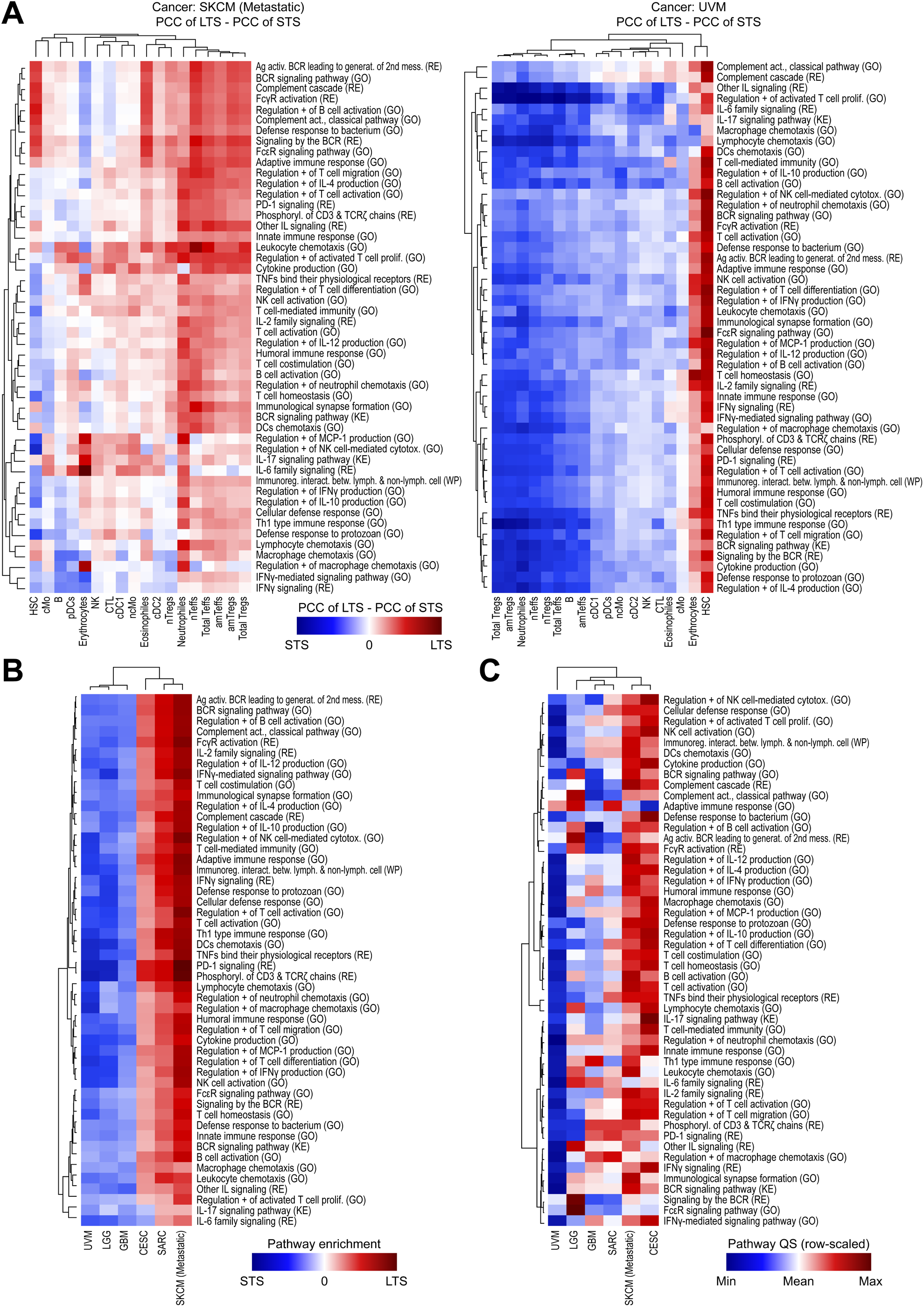
Altered LTS and STS differential correlations between immune pathways and cell signatures in immune-privileged cancer sites versus immunogenic cancers. (A) Heatmap representations of the difference between LTS minus STS PCC values from the Figure 4 for immunogenic cancer SKCM (Metastatic) (left) and immune-privileged cancer UVM (right). Negative values are going from blue to white and denote a stronger association with STS, whereas positive values are going from white to red and denote a stronger association with LTS. Dendrogram clustering were realized for both rows and columns. (B) Heatmap illustrating raw pathways enrichment values for immune-related pathways in the six previously depicted cancers (three immunogenic: CESC, SARC and SKCM (Metastatic), and 3 immune-privileged: UVM, GBM and LGG). Negative values are going from blue to white and denote a stronger pathway enrichment in STS, whereas positive values are going from white to red and denote a stronger pathway enrichment in LTS. Dendrogram clustering were realized for both rows and columns. (C) Row-scaled heatmap showing the whole patients averaged QS computed for each depicted immune-related pathway in the six previously mentioned cancers. Minimum row values are going from blue to white whereas maximum row values are going from white to red. Dendrogram clustering were realized for both rows and columns.

Indeed, profiling pathway enrichment raw values alone in the LTS and STS groups of the six cancers reflected their effect on survival (**Figure 5B**). In the three immunogenic cancers SKCM (Metastatic), SARC, and CESC, almost all the immune-related pathways are significantly enriched (enrichment values > 0 and adjusted p-values < 0.05) in LTS as compared to STS, whereas they are effectively “enriched” (values < 0 and adjusted p-values < 0.05) in STS in the three immune-privileged cancers, GBM, LGG and UVM. Thus, we confirmed that the same exact immune pathways confer good or bad prognosis dependent on a cancer origin.

We reasoned that, if pathway enrichment analysis does not discriminate between pathways that confer good or bad prognosis, then looking at the abundance (QS) of the pathways, without using any pathway enrichment metric nor survival information, may provide clues about the underlying differences between tumors from immune-privileged vs. those from non-immune-privileged tissues. Indeed, we found that the immune-related pathways tend to be expressed at a lower level in UVM, LGG and GBM, whereas they show higher relative expression in immunogenic cancers BRCA, SARC and SKCM (Metastatic) (**Figure 5C**). In the light of the pervasive evidence that any immune activity is harmful to patients with UVM, LGG or GBM, this last result means that even a minimal immune system activity confers a negative effect on the survival of patients with cancer of immune-privileged site.

Moreover, the integration of the findings in the *AMOCATI*-defined STS and LTS subsets with the “outlier” QS detected by the pathway abundance analysis may indicate which pathways may be a promising target for treatment in a given cancer. For instance, in the SARC, the *Th1 type immune response* pathway is significantly more enriched in LTS than in STS, making it of good prognosis (**Figure 5B**). But the expression of this pathway in relatively low in SARC, as compared to other immunogenic cancers (**Figure 5C**), which may constitute a potential target for activation in this cancer type.

Taken together, *AMOCATI*-based analysis of different types of cancer revealed important differences in the associations between biological pathways and individual immune subsets, pointing out pathological nature of the immune-related functions that may be responsible for poor outcomes in patients with LGG, UVM or other tumors of the immune-privileged sites.

### DISCUSSION AND PERSPECTIVES

Our new analysis workflow, *AMOCATI*, is a powerful computational tool to identify exceptional survival responses in cancer patient cohorts based only on transcriptomic data and survival information. Here, we use this power to find meaningful associations between immune-infiltrating cells and immune-related pathways with survival in the transcriptomic datasets of patients with different types of cancer. We find that some tumor transcriptomes contain robust immune cell signatures, while in others the transcriptional evidence of immune infiltration is low. However, the overall immune infiltration within tumors can have a positive or a negative impact on patient survival depending on cancer type and location. Importantly, using *AMOCATI*, we are able to show that high or low infiltration *per se* is not a good or bad prognostic marker. Some cancers are well infiltrated by the immune system, but this infiltration is associated with shorter survival. Some other cancers are poorly infiltrated, but even a modest increase in infiltration is associated with a poor outcome, such as in cancers of immune-privileged organs. By using transcriptional data from short- and long-term survivors classified by *AMOCATI*, we obtained a refined picture of tumor-immune system interactions, and linked poor outcomes in cancers of immune-privileged sites to the infiltration by the immune system.

Our results support and extend recent works that use computational approaches to study tumor-immune system interactions. We confirmed that a large-scale transcriptomics analysis can identify two classes of tumors relative to immune orientation: those, in which activation of immune pathways is generally associated with a better outcome, and those in which this activation has a negative effect on survival(5, 6, 9, 10). However, there has been no consensus regarding the biological causes of this observation. Despite using different approaches for defining the tumor immune subtypes, these studies reported substantial variation in the proportion of the immune subtypes in the individual tumor types and TCGA subtypes, painting an overwhelmingly complex picture of tumor-immune system interactions. Although almost all studies noted that patients with brain and/or eye tumors fare worse when there is evidence of immune infiltration, some noted that tumors of the eye and/or brain cluster in a separate immune subtype(5, 10), where the immune orientation was not linked to cancer origin. We chose an agnostic approach with regards to the possible molecular or mutational profiles within clinician-defined cancer types. By considering solely the pathological diagnosis and the length of survival, we let *AMOCATI* guide the choice of transcriptional signatures associated with a better or worse outcome, and scrutinized these signatures for evidence of immune system activity. As a result, a clear picture emerged: many cancers in which immune infiltration has a negative effect on survival arise in the immune-privileged tissues, such as the eye (uveal melanoma), the brain (gliomas and glioblastomas) or the testis (testicular germ cell tumor).

In this regard, our study reproduces several isolated observations that in patients with low-grade glioma, glioblastoma, or uveal melanoma, the clinical outcome is compromised when there is transcriptional evidence of the immune system involvement(48–51). By looking at transcriptional profiles of all of these tumor types in parallel, we found a consistent pattern: a low overall infiltration, which suggests an immune-protected status, with evidence of shorter survival following even a minimal increase in infiltration by classical anti-cancer players (NK cells, cMo, ncMo, cDC1, cDC2 and B cells). The same pattern was also seen in TGCT. We think that an underlying cause of this observation is the immune-privileged status of the tissue in which these tumors arise (eye, brain, and testis). Whether the poor outcome is a consequence of this infiltration (for example due to growth promoting effects of the inflammatory cytokines on tumor cells), or a sign of the disruption of the immunological barrier during tumor invasion, remains to be tested. What is clear is that the immune subsets and pathways linked to poor survival in these tumor types are similar, if not identical, to those that improve survival in other cancer types. Therefore, classical immunotherapy approaches, aimed at disinhibiting the anticancer-immune response, are to be avoided in patients with tumors of the immune-privileged sites. The clinical experience supports this recommendation(52–54).

In conclusion, *AMOCATI* is a new tool for studying large transcriptional datasets. The advantage of this method is that it classifies patients based on their transcriptional signatures alone, and needs only minimal clinical information, such as clinical diagnosis and duration of survival, to study the association of transcripts with survival. This extends the range of data that can potentially be compiled for *AMOCATI*-based study, which may be especially useful for rare conditions. We were able to harness this power to uncover the deleterious effect of immune infiltration and immune activity in tumors from immune-privileged sites, confirming at least in a set of cancers the 160 years-old theory of Rudolph Virchow that immune infiltrates may facilitate cancer progression(12). We hope that *AMOCATI* would be of use in studies of other diseases and in other experimental settings.

## ACKNOWLEDGMENTS

We thank Damien Chaussabel for discussion. The human results reported here exclusively used data generated by the TCGA/TARGET Research Networks: https://www.cancer. gov/ccg/research/genome-sequencing.

## AUTHOR CONTRIBUTIONS

G.D.-J. conceived the study. GDJ and PR. conceived the principle of *AMOCATI*. P.R. elaborated and wrote the AMOCATI package script. P.R. performed the analysis. P.R., G.D.-J. and K.P analyzed and interpreted the data. N.C. validated the approach. P.R., G.D.-J. and K.P. wrote the manuscript.

## DATA AND MATERIAL AVAILABILITY

*Human cancer RNA-Seq data*: survival and RNASeq data from the TCGA cancer project were freely downloaded from the GDC Cancer Portal of the National Cancer Institute (USA) (https://portal.gdc.cancer.gov/) (Files from release versions 11 and 12).

*Generation of cell subsets genes signatures*: the following datasets accession numbers (NCBI GEO) were used to generate custom gene signatures for immune cell subsets: GSE94820, GSE35457, GSE28490, GSE28491, GSE89232, GSE6887, GSE60448, GSE24759, GSE65010 and GSE76598. See also **Table S1**.

## ABBREVIATIONS

ACC: adrenocortical carcinoma
ALL-P2: acute lymphoblastic leukemia, phase II
ALL-P3: acute lymphoblastic leukemia, phase III
AML: acute myeloid leukemia
amT_effs_: activated/memory CD4^+^ T effector cells
amT_regs_: activated/memory CD4^+^ T regulatory cells
BLCA: bladder urothelial carcinoma
BLGSP: Burkitt lymphoma genome sequencing project
BRCA: breast invasive carcinoma
BM: bone marrow
CESC: cervical squamous cell carcinoma and endocervical adenocarcinoma
CHOL: cholangiocarcinoma
cDCs: Flt3^+^ Zbtb46^+^ conventional dendritic cells
cDC1: CD141^+^ cDCs
cDC2: CD1c^+^ cDCs
cMo: CD14^++^CD16^neg^ classical monocytes
COAD: colon adenocarcinoma
CS: clinical score
CTL: cytotoxic CD8^+^ T lymphocytes
DLBC: lymphoid neoplasm diffuse large B-cell lymphoma
ECM: extracellular matrix
ESCA: esophageal carcinoma
FPR: false positive rate
GAG: glycosaminoglycan
GBM: glioblastoma multiforme
GO: gene ontology
HNSC: head and neck squamous cell carcinoma
HSC: hematopoietic stem cells
HTMCP-CC: HIV^+^ tumor molecular characterization project (cervical cancer)
IS: immune system
KE: KEGG (Kyoto Encyclopedia of Genes and Genomes)
KICH: kidney chromophobe
KIRC: kidney renal clear cell carcinoma
KIRP: kidney renal papillary cell carcinoma
LAML: acute myeloid leukemia
LGG: brain lower grade glioma
LIHC: liver hepatocellular carcinoma
LTS: long-term survivors
LUAD: lung adenocarcinoma
LUSC: lung squamous cell carcinoma
MESO: mesothelioma
NBL: neuroblastoma
ncMo: CD14^+^CD16^++^ nonclassical monocytes
NK: natural killer cells
nT_effs_: naïve CD4^+^ T effector cells
nT_regs_: naïve CD4^+^ T regulatory cells
OS: osteosarcoma
OV: ovarian serous cystadenocarcinoma
PAAD: pancreatic adenocarcinoma
PB: peripheral blood
PCPG: pheochromocytoma and paraganglioma
pDCs: plasmacytoid dendritic cells
PRAD: prostate adenocarcinoma
QS: quantitative score
READ: rectum adenocarcinoma
ROC: receiver operating characteristic
SARC: sarcoma
SD: standard deviation
SKCM: skin cutaneous melanoma
STAD: stomach adenocarcinoma
STS: short-term survivors
TGCT: testicular germ cell tumors
THCA: thyroid carcinoma
THYM: thymoma
TPR: true positive rate
UCEC: uterine corpus endometrial carcinoma
UCS: uterine carcinosarcoma
UVM: uveal melanoma
WP: WikiPathways
WT: high-risk Wilms tumor

