## Supplementary material for "Application of the *AMOCATI* R workflow to tumor transcriptomic data delineates the adverse effect of immune cell infiltration in immune-privileged organs": Figure S1

### Supplemental Figure 1

A

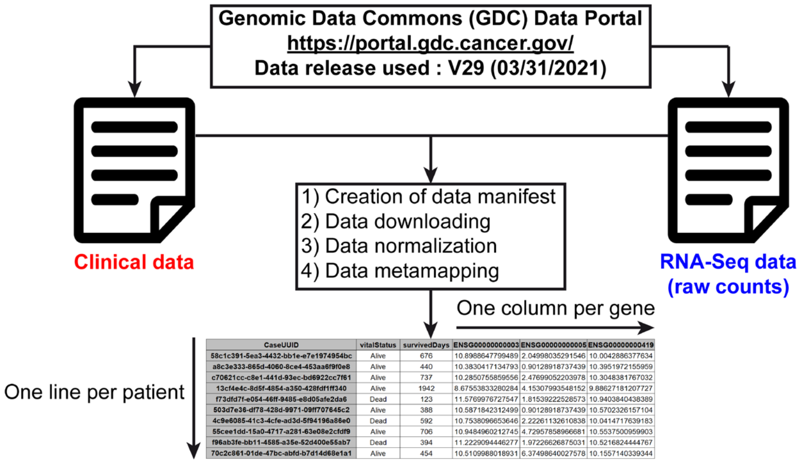


B

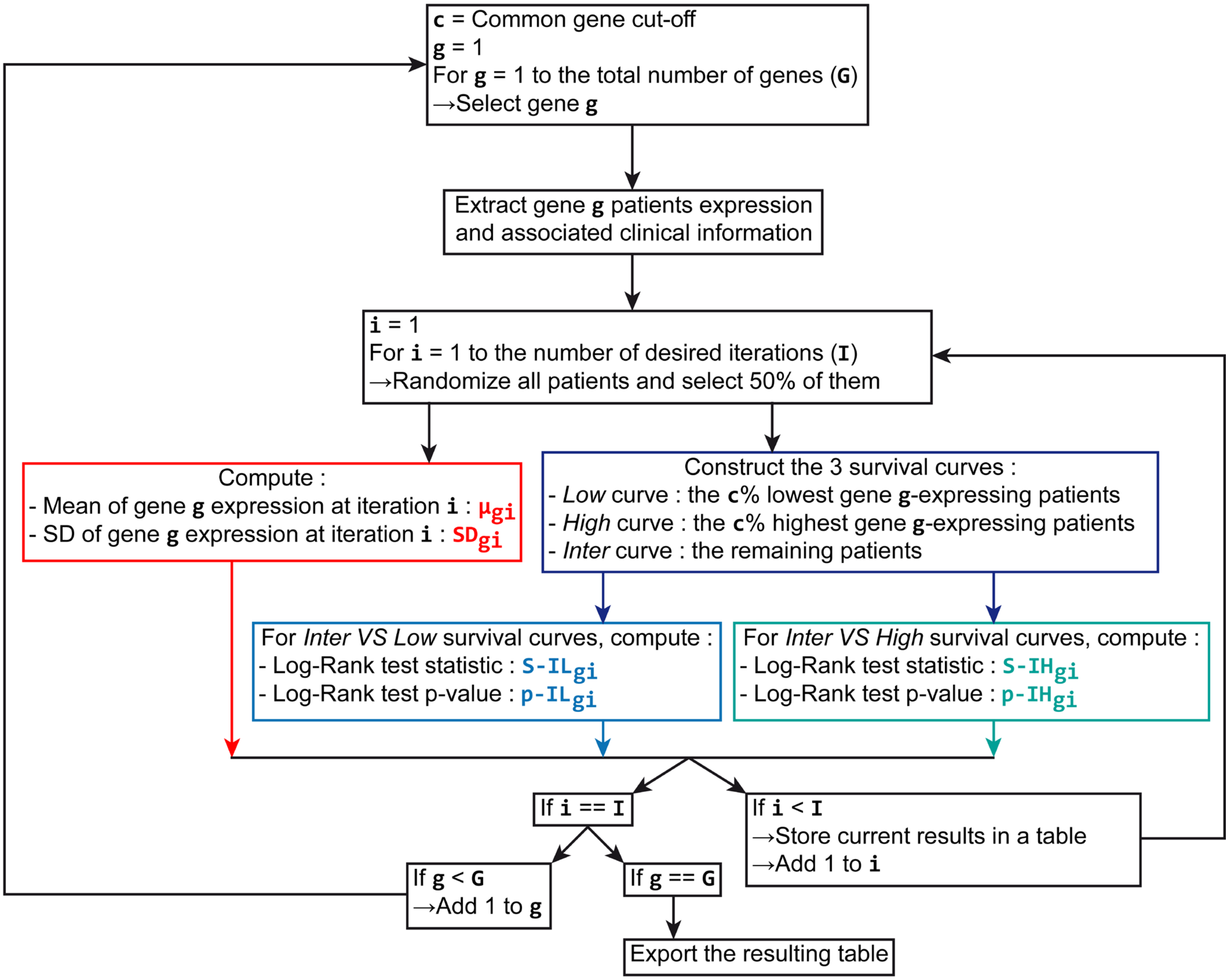


**Figure S1 (part 1)**

**C**

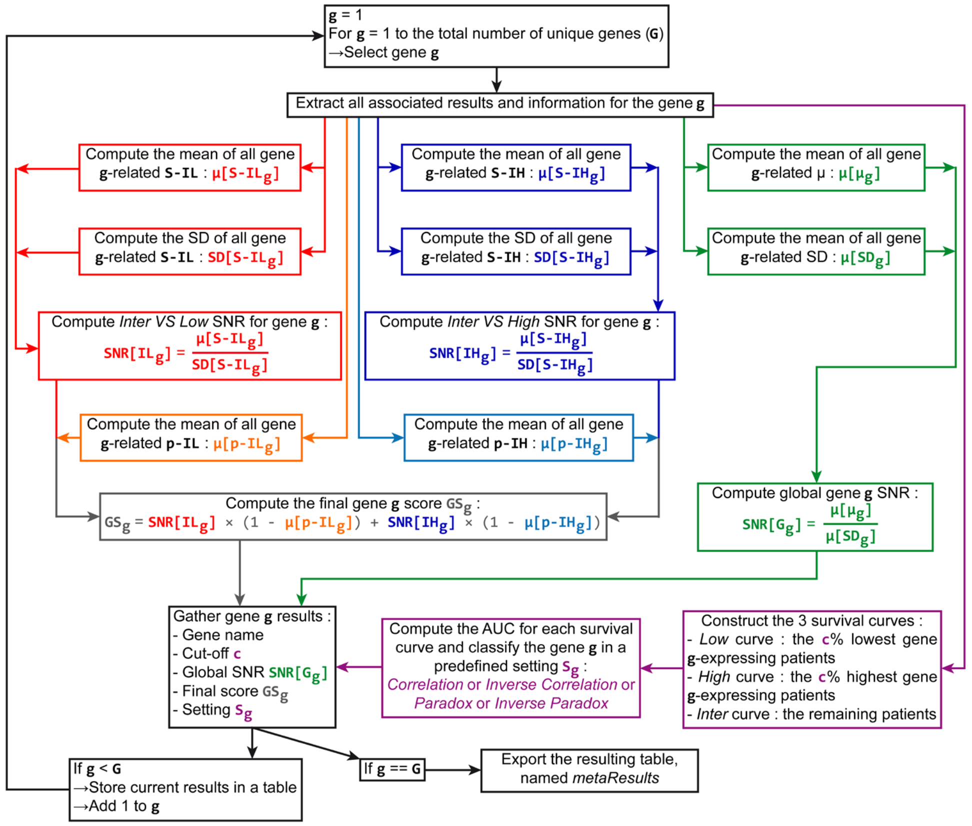


**D**

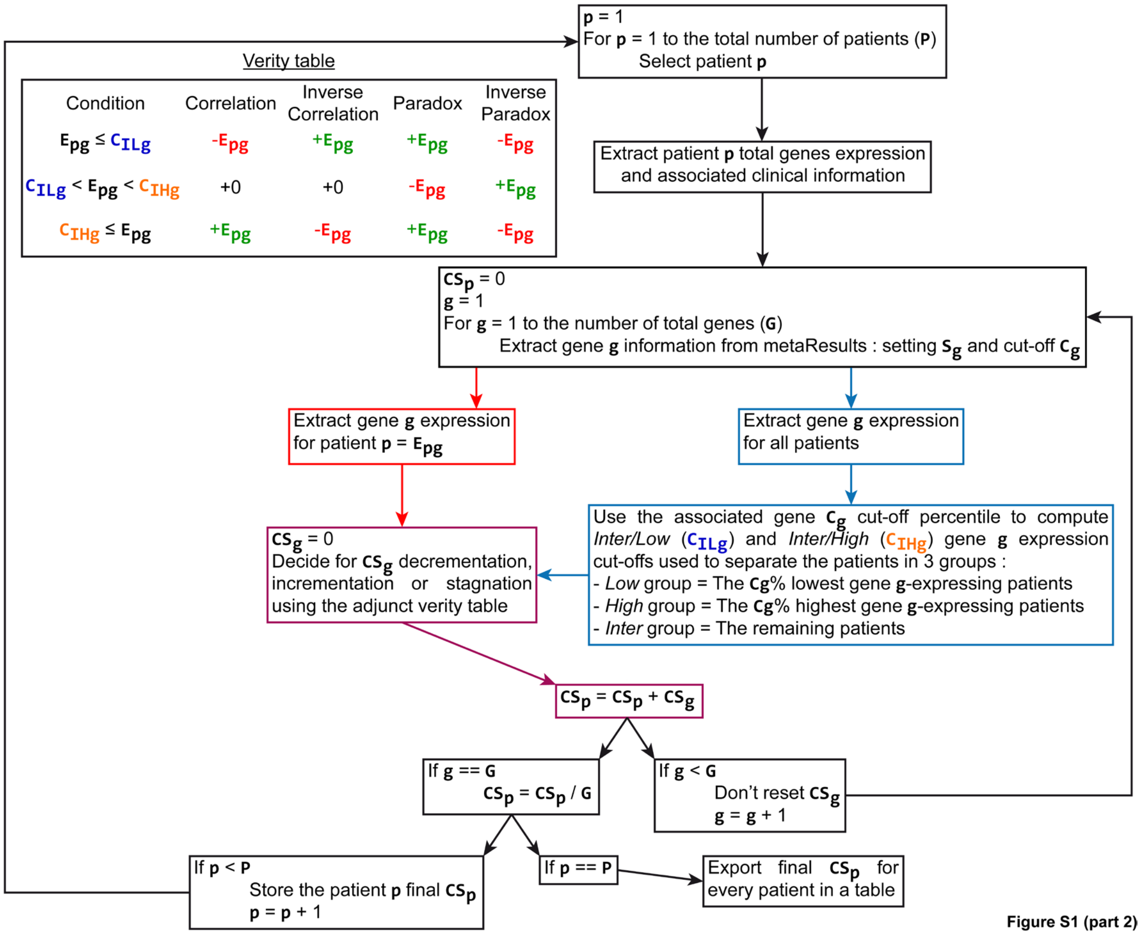


Verity table

| Condition | Correlation | Inverse Correlation | Paradox | Inverse Paradox |
|---|---|---|---|---|
| $E_{pg} \leq C_{ILg}$ | $-E_{pg}$ | $+E_{pg}$ | $+E_{pg}$ | $-E_{pg}$ |
| $C_{ILg} < E_{pg} < C_{IHg}$ | +0 | +0 | $-E_{pg}$ | $+E_{pg}$ |
| $C_{IHg} \leq E_{pg}$ | $+E_{pg}$ | $-E_{pg}$ | $+E_{pg}$ | $-E_{pg}$ |

**Figure S1 (part 2)**

**Figure S1 – Generation of an *all-in-one* file and presentation of the *metaResults* and *Clinical Score* computation methods. (A)** UML diagram describing the process used for the generation of *all-in-one* data files for a given cancer project from GDC repository. **(B)** UML diagram describing the bootstrapping algorithm (first part of the *metaResults* method) used to estimate iteratively the impact of each gene on low-, intermediate- and high-expressing patients survival. **(C)** UML diagram describing the second part of the *metaResults* method, used to synthesize and generate the final *metaResults* file for a given dataset using the bootstrapping results obtained in **(B)**. **(D)** UML diagram describing the algorithm used to compute the *Clinical Score* of each patient for a given gene signature.
