## Supplementary material for "Application of the *AMOCATI* R workflow to tumor transcriptomic data delineates the adverse effect of immune cell infiltration in immune-privileged organs": Figure S2

### Supplemental Figure 2

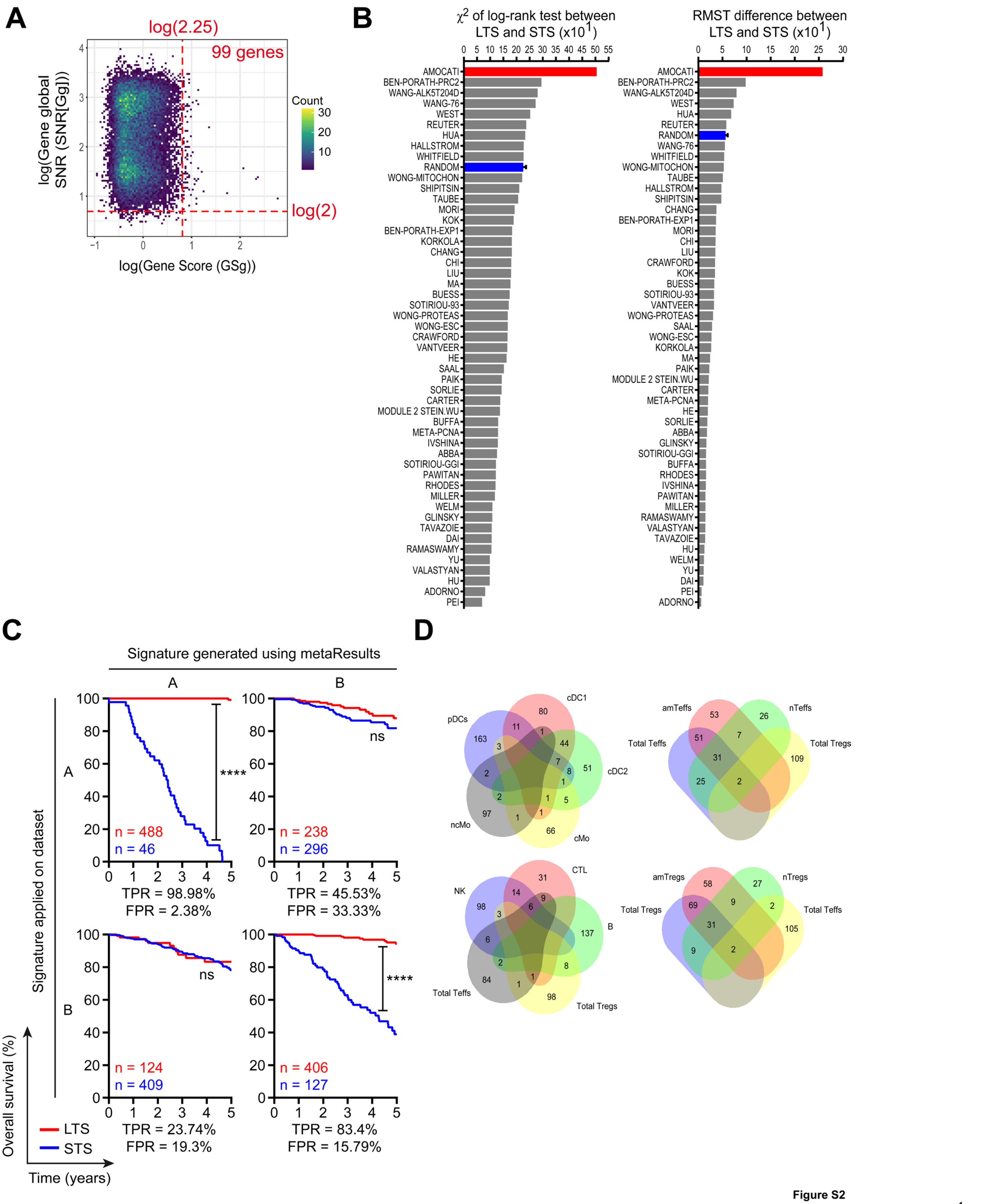


**Figure S2**

**Figure S2 – Use of the AMOCATI workflow to isolate and evaluate signatures in BRCA cancer and Venn diagram of our cell subset-specific gene signatures. (A)** Dot plot showing each gene Gene Score ($GS_g$) and global SNR ($SNR[G_g]$) as well as their respective thresholds used to isolate the final classification signature (upper-right quarter) for the BRCA cancer. **(B)** Bar plot showing the efficiency of *AMOCATI*-determined classification signature to separate LTS and STS in BRCA cancer (red) as compared to ten randomly-generated signatures (blue) and to previously published prognosis signatures by several authors and teams (grey). **(C)** Kaplan-Meyer plots showing in BRCA the impossibility of prediction uses of *AMOCATI*'s determined classification signature. BRCA patients were randomly split into two equal subdatasets A and B, and respective *metaResults* and classification signatures were determined independently for each. Then, each classification signature was applied on each dataset using its respective *metaResults* file, leading to the 4 different combinations presented here. **(D)** Venn diagrams showing the overlap between the final gene signatures obtained using publicly available cell-specific GEO datasets. Myeloid cell-related gene signatures (pDCs, cDC1, cDC2, ncMo and cMo) are shown in the top-left diagram and lymphoid cell-related gene signatures (NK, CTL, B-cells, total Tregs and total Teffs) in the bottom-left one. Further insight about $T_{effs}$ and $T_{regs}$ subsets are given in the top-right and bottom-right diagrams, respectively. Numbers inside each case indicate the number of genes that are common between the depicted signatures.
