## Supplementary material for "Application of the *AMOCATI* R workflow to tumor transcriptomic data delineates the adverse effect of immune cell infiltration in immune-privileged organs": Figure S3

### Supplemental Figure 3

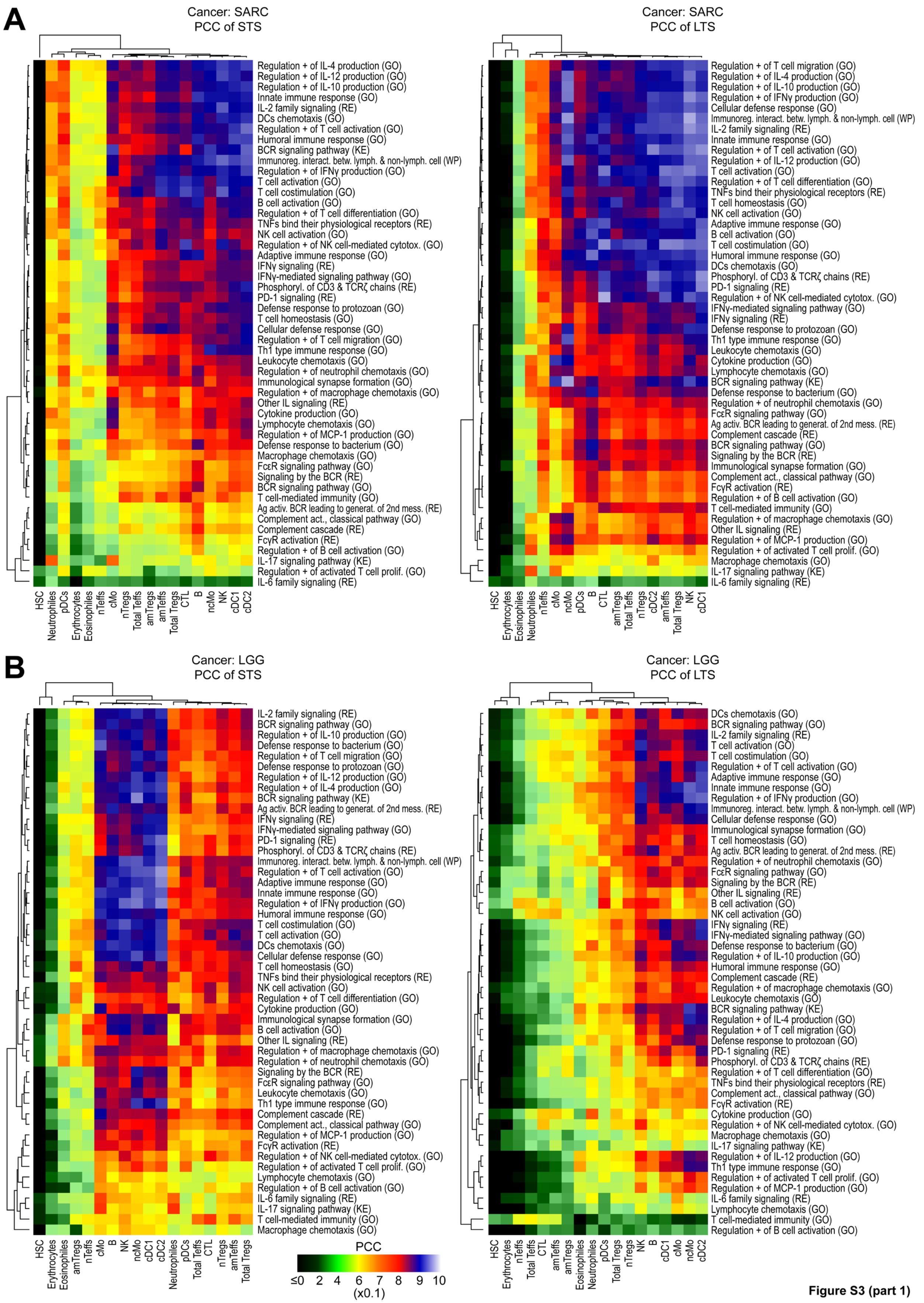


**Figure S3 (part 1)**

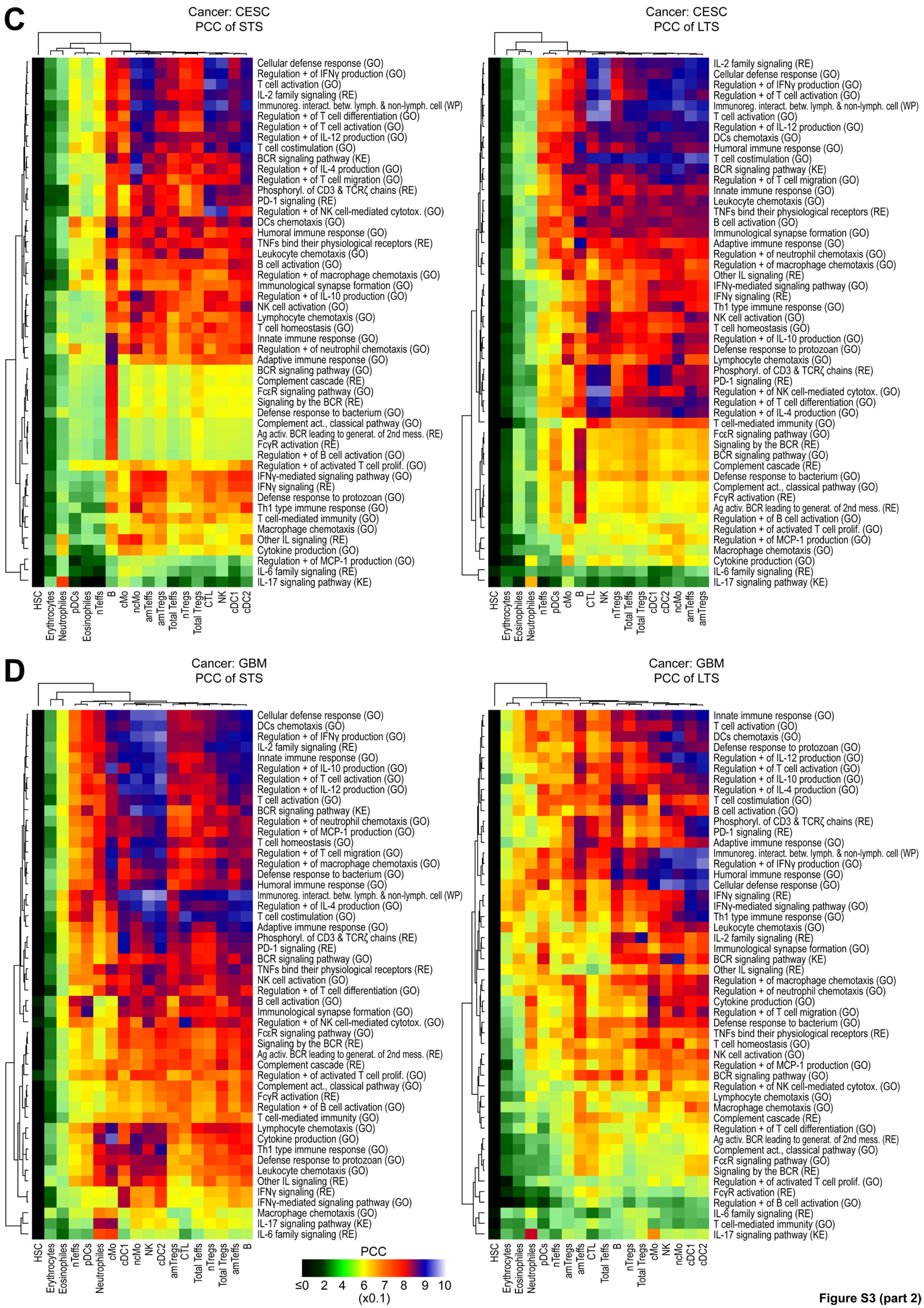


**Figure S3 (part 2)**

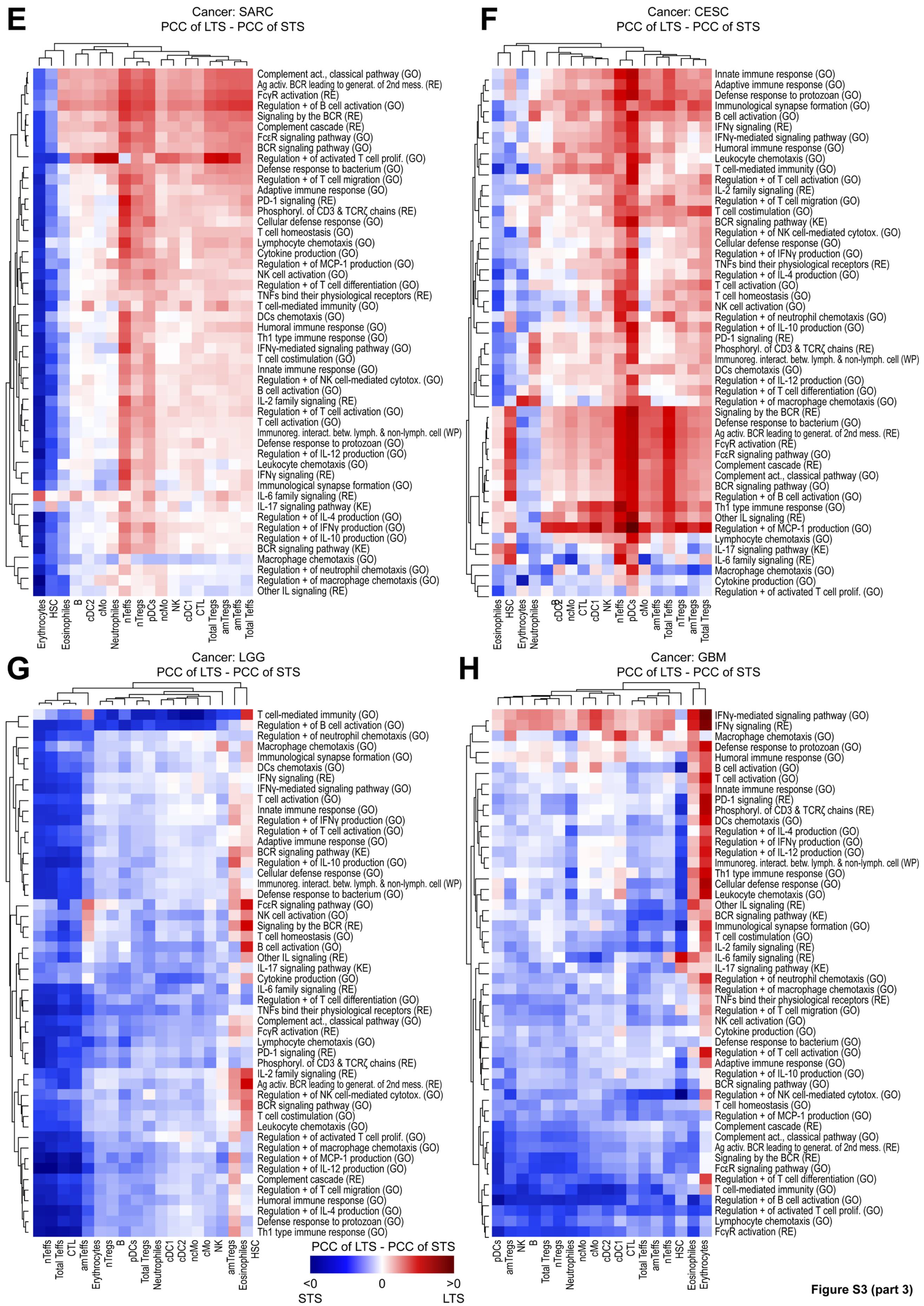


**Figure S3 (part 3)**

**Figure S3 – Altered correlations between pathways and cell populations signatures in SARC, LGG, CESC and GBM cancer biopsies. (A-D)** Heatmaps showing the PCCs obtained for each pathway and cell population in two immunogenic cancers (SARC in **(A)** and CESC in **(C)** and in two immune-privileged cancers (LGG in **(B)** and GBM in **(D)**) and. For each subfigure, STS and LTS patients are represented separately, with STS heatmap on the left, and LTS heatmap on the right. Dendrogram clustering were realized for both rows and columns. Black to yellow colors represent PCCs between 0 and 0.6, yellow to orange between 0.6 and 0.7, orange to red between 0.7 and 0.8, red to blue between 0.8 and 0.9, and blue to white between 0.9 and 1. **(E-H)** Heatmaps showing the PCCs difference (LTS minus STS) for each depicted immune-related pathway and cell population in two immunogenic cancers (SARC in **(E)** and CESC in **(F)**) and in two immune-privileged cancers (LGG in **(G)** and GBM in **(H)**). Negative values are going from blue to white and denote a stronger correlation in STS, whereas positive values are going from white to red and denote a stronger correlation in LTS. Dendrogram clustering were realized for both rows and columns.
